# Non-null Effects of the Null Range in Biogeographic Models: Exploring Parameter Estimation in the DEC Model

**DOI:** 10.1101/026914

**Authors:** Kathryn A. Massana, Jeremy M. Beaulieu, Nicholas J. Matzke, Brian C. O’Meara

## Abstract

Historical biogeography seeks to understand the distribution of biodiversity in space and time. The dispersal-extinction-cladogenesis (DEC) model, a likelihood-based model of geographic range evolution, is widely used in assessing the biogeography of clades. Robust inference of dispersal and local extinction parameters is crucial for biogeographic inference, and yet a major caveat to its use is that the DEC model severely underestimates local extinction. We suggest that this is mainly due to the way in which the model is constructed to allow observed species to transition into being present in no areas (i.e., null range). By prohibiting transitions into the null range in the transition rate matrix, we were able to better infer local extinction and support this with simulations. This modified model, DEC*, has higher model fit and model adequacy than DEC, suggesting this modification should be considered for DEC and other models of geographic range evolution.

## Introduction

Historical biogeography has developed from simply observing the general patterns of species, to incorporating events that explain biogeographic processes (such as vicariance and dispersal), to developing explicit probabilistic approaches. With the advent of parametric methods based on maximum likelihood and Bayesian frameworks, researchers have been able to incorporate important information, such as branch lengths and fossils (e.g., allowing for better tree dating estimates) (Smith and Donoghue 2010; Wood et al. 2012; Beaulieu et al. 2013a; Pyron 2014).

A popular method (cited 499 times since its publication) to assess the historical biogeography of taxa is the dispersal-extinction-cladogenesis (DEC) model (Ree and Smith 2008), which estimates geographic range evolution for anagenetic (i.e., along branches) and cladogenetic (i.e., at nodes) change on a phylogeny. In the case of anagenetic change, range expansion and range contraction [modeled as parameters by way of the rate of dispersal from area *i* to area *j* (D_ij_) and local extinction in area *i* (E_i_)] are modeled as stochastic processes along branches. Published analyses with DEC assume the simplest model involving only a single dispersal rate and a single local extinction event, although it is possible to have more complex models (i.e., allowing different dispersal rates among each area pair by having *n*^2^*-n* free parameters for the dispersal rates, and *n* free parameters for local extinction). A rate matrix can be assembled for a given number of geographic ranges and rate parameters (Fig. 1).

**Figure 1.**
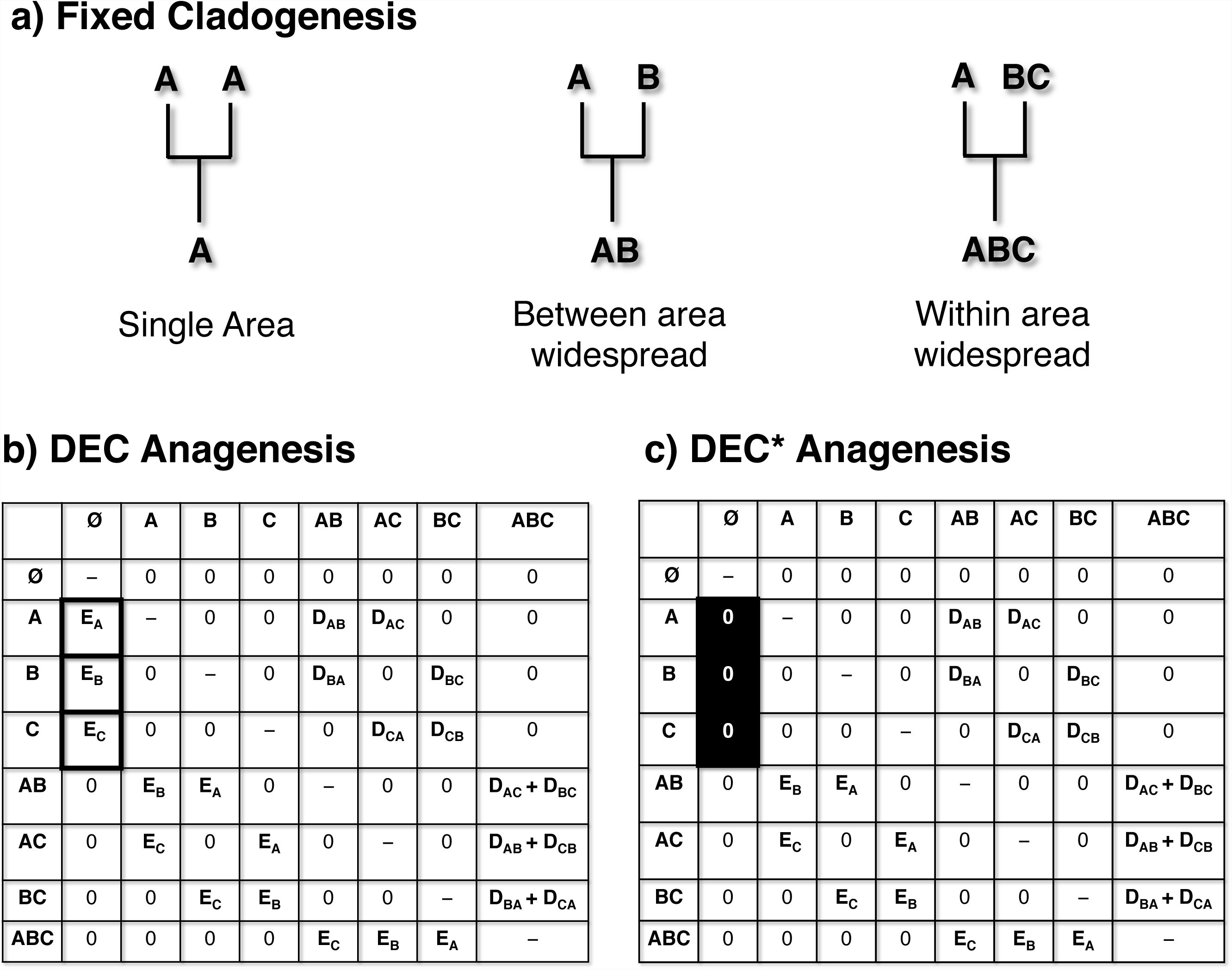
Diagram showing (a) the allowed cladogenetic events for DEC and DEC*, (b) the anagenesis transition rate matrix for DEC, and (c) the DEC* anagenesis transition rate matrix, assuming three geographic states (A, B, C). At a cladogenetic event, if a species is in one area, the descendant species inherits that area. If the species is in multiple areas, one species inherits one area, while the other species inherits all areas (peripatric speciation; within area widespread) or it is allowed to inherit all areas but the area occupied by the first species (vicariant allopatric speciation; between area widespread). Note that extinction is not allowed in DEC* from one state to zero states. D = dispersal, E = local extinction.

Ree and Smith (2008) carried out simulations to test the accuracy of the DEC model on parameter estimation. They found that although the model worked reasonably well, dispersal was underestimated and local extinction was severely underestimated, often estimated as being effectively zero. Note that while there has been a robust discussion of whether extinction rates can be estimated on molecular phylogenies (Nee et al. 1994; Rabosky 2010; Beaulieu and O’Meara 2015), the extinction rate in that case relates to speciation and extinction of entire species and its signal on a phylogeny’s topology and branch lengths. The extinction rate relevant for our purposes is the rate of having a population no longer occurring in a particular area, using a fixed tree. In terms of its role in a model, it is more like the rate of reduction in a meristic character than a rate at which a species goes extinct, even though biologically it appears more similar to the latter. Thus, its difficulty in being estimated is surprising.

One feature of the DEC model that has received little comment is that it includes a null range (a geographic range of 0 areas) in the anagenesis transition matrix (Fig. 1). In one sense, inclusion of the null range is a natural modeling decision, since the assumption that local extirpation is a process directly implies that the same process can reduce a single area geographic range to a range of size 0. However, the inclusion of a null range in the state space has some peculiar properties. For instance, no sampled species will ever occupy the null range state; even extinct species, if included in an analysis, are included because they occurred in some area. We suspect that the only way to fit any data pattern that does not observe null ranges is by driving down the rate of range contraction to the point where the probability of such an event is effectively zero. Unfortunately, given that all transitions to other extinction scenarios are linked through a global extinction parameter it seems unavoidable that when null ranges are allowed in the model extinction would generally be underestimated.

Of course, in some ways, the extinction rate is a nuisance parameter – that is, the hundreds of studies using the DEC model primarily focus on ancestral state estimates rather than on rates. However, given that this rate represents one of the two free parameters that are then used for inferring ancestral states on the tree, we expect that biased extinction estimates may result in errors in the ancestral state estimation. In other words, given low extinction rate estimates, areas can only be lost at speciation events, so we predict a greater number of areas as we move rootward on the tree and few area losses along longer branches. This study attempts to modify the DEC model to improve estimation of extinction rate and then tests using simulated and empirical data to see if this results in a better model overall.

## Methods

We modified the DEC model (which we refer to DEC* hereafter) to omit transitions into the null state in the anagenesis transition rate matrix between ancestor and descendent pairs (Fig. 1). It has the same number of parameters as DEC (dispersal and local extinction, *d* and *e* respectively), with the only change being fixing the transition rate to 0 for transitions from ranges of size 1 to the null range. DEC* is distinct from the three-parameter DEC+J model which allows for founder-event speciation associated with lineage-splitting with the addition of the free *j* parameter (Matzke 2014b). DEC+J retains the DEC assumption that a null geographic range is a valid state. To implement the DEC* model, we modified the original lagrange DEC C++ code (https://github.com/rhr/lagrange-cpp) to omit the transition into the null range in the anagenesis transition rate matrix. The DEC* model is implemented as a modification to the DEC C++ version, and is also allowed in the BioGeoBEARS R package (Matzke 2013) by changing the *include_null_range* setting from TRUE to FALSE in the BioGeoBEARS model setup.

We implemented our own DEC simulator in R that follows the procedures described by Ree and Smith (2008). The simulator produces birth-death phylogenetic trees with concurrent range evolution, combining the DEC model and stochastic cladogenesis. The simulator also does the same for the DEC* model. Trees were produced with the same known dispersal and local extinction parameter, constrained to vary between 0.01 and 0.2, while speciation was constrained to be 0.4 events per million years. Geography consisted of three possible hard-coded geographic areas, meaning that there were 8 possible geographic ranges in the state space of the DEC simulation (A, B, C, AB, AC, BC, ABC, and null), and 7 possible ranges in the DEC* simulation (as the null range state is excluded). At cladogenesis, when the lineage had a widespread range, equal probabilities were assigned to each allowed range-inheritance scenario (vicariance or subset sympatry). For both DEC and DEC*, we performed 2,000 simulation-inference runs and compared dispersal and local extinction parameter estimates as well as the number of correctly inferred number of areas at internal nodes for all simulations. The simulations began by assigning the root node a range of a random single geographic area. The phylogeny was allowed to grow according to the DEC or the DEC* model until it reached 100 taxa (extant plus extinct). To match empirical datasets, the simulated phylogenies were pruned of branches that went extinct.

Our main objective was to understand DEC* versus DEC analyses on empirical datasets. Therefore, we searched the literature for published studies that used the DEC model. Then we compiled the phylogenies and geography presence/absence data available, which resulted in 15 empirical datasets. Most of these datasets were used in Matzke (2014b), and followed any modifications made therein (Kambysellis et al. 1995; Baldwin and Sanderson 1998; Hormiga et al. 2003; Jordan et al. 2003; Clark et al. 2008; Dunbar-Co 2008; Ree and Smith 2008; Benavides et al. 2009; Clark et al. 2009; Gillespie and Baldwin 2010; Smith and Donoghue 2010; Lerner et al. 2011; Nicholson et al. 2012; Bennett et al. 2013; Lapoint et al. 2013; Matzke 2014a). However, we also assessed the caecilian and salamander datasets from a recent published study using DEC+J (Pyron 2014), and a palpimanoid spider dataset (Wood et al. 2012). We performed unconstrained analyses with C++ DEC and DEC* on each dataset. We compared analyses between DEC, DEC*, and DEC+J for all 15 datasets. If the values were not available, we used the package BioGeoBEARS (Matzke 2013) to run all DEC+J analyses. We also assessed model adequacy for each dataset by comparing the number of areas estimated per node between DEC and DEC* to the observed modern geographic range sizes.

## Results

### Simulations

Figure 2 shows the observed parameter estimates of local extinction and dispersal for DEC and DEC* compared to the true estimates under a DEC simulation. Overall, the point estimate for local extinction was closer to the true value under DEC* than with DEC (Fig. 2), although with higher variance. With simulations under the DEC model we found that the median for local extinction under a DEC* inference (*e*=0.0957) was closer to the true local extinction median estimate (*e*=0.0989), while the median for local extinction under DEC inference was close to zero (*e*=1.287e-06). Similarly, with simulations under the DEC* model we found that the median local extinction estimated under DEC* inference (*e*=0.1028) is almost identical to the true local extinction parameter (*e*=0.1030), whereas, again, local extinction is grossly underestimated under the DEC inference (*e*=1.200e-6).

**Figure 2.**
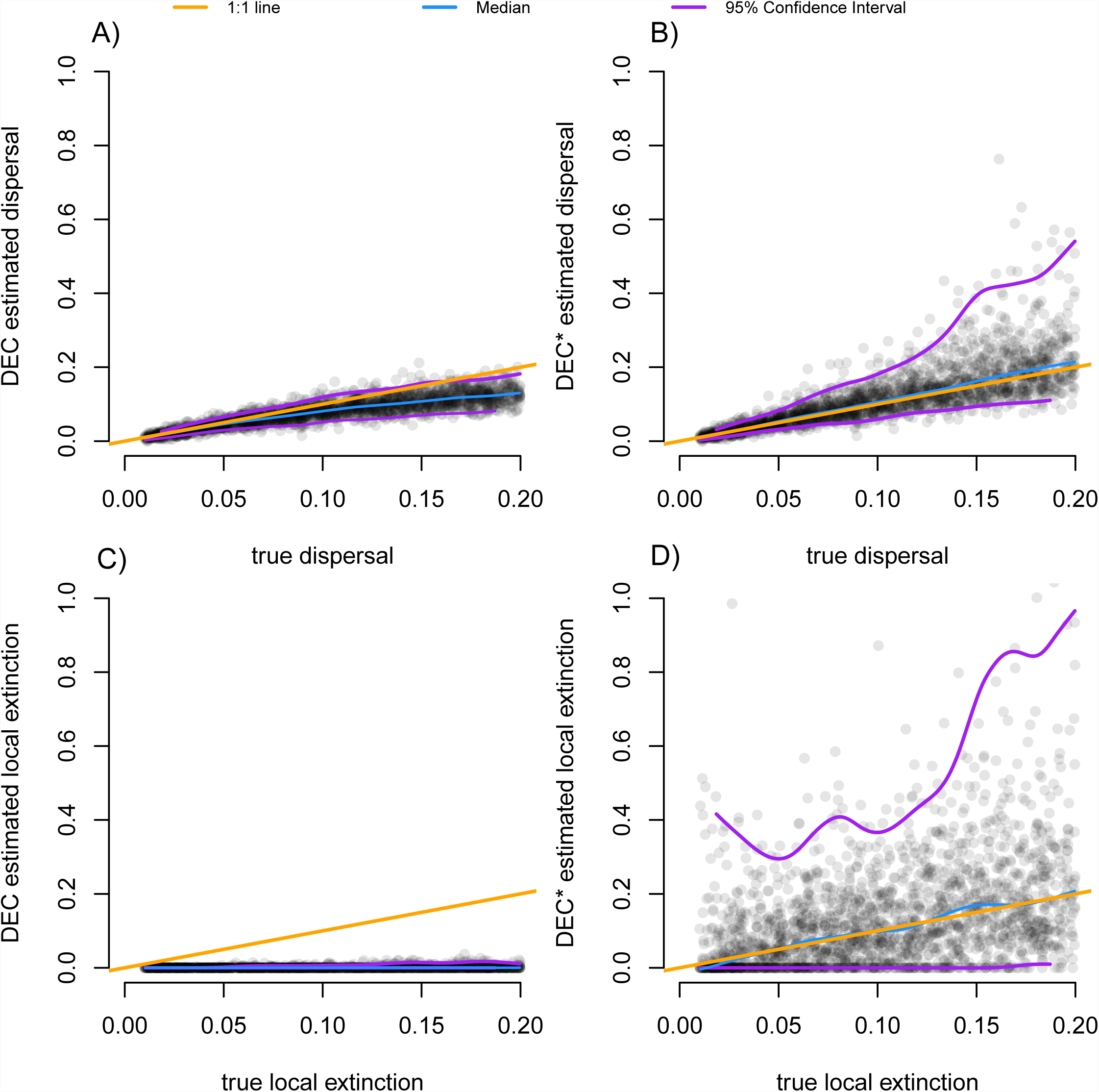
Plots showing parameter inference under the 2,000 DEC simulations. DEC inference of dispersal (A) was not as effective as dispersal inferred under DEC* (B). Local extinction under DEC inference (C) was highly underestimated. Local extinction under DEC* (D) was better estimated in comparison to DEC inference, although with more variance. Purple lines represent 95% confidence intervals; blue line shows the median; orange line shows the 1:1 line.

Median estimates of dispersal under DEC simulations were closer to the median of the true dispersal parameter (*d*=0.0989) under DEC* than under DEC inference (*d*=0.0766) (Fig. 2). When simulating under DEC* (Supplemental Fig. 1), the median dispersal under DEC* inference (*d=0*.1053) was again closer to the median dispersal rate used in the simulation (*d*=0.1030) than dispersal inferred under DEC (*d*=0.0789).

We calculated the root mean square error (RMSE) of the estimated parameter values for DEC and DEC*. The root mean square error gives the standard deviation associated with the differences between the true parameter and the inferred parameter estimates, and here a smaller value indicates less error in the parameter inference. Results indicated that on the logarithmic scale the error for *e* was far better for DEC* than DEC and nearly the same for *d* (RMSE of *e* was 10.9120 for DEC and 3.8363 for DEC*; RMSE of *d* was 0.3784 for DEC and 0.3886 for DEC*). However, on a linear scale, error is far better for both parameters for DEC than DEC*, due to some tremendously high values of *e*. (RMSE of *e* was 0.1135 for DEC and 2.2345 for DEC*; RMSE of *d* was 0.0363 for DEC and 0.7816 for DEC*).

Finally, we assessed the accuracy of DEC against DEC* in estimating the geographic area range at the root. Under DEC simulation, the root state was correctly estimated 49.05% of the time, whereas under DEC*, 58.30% of root states were accurately estimated.

### Empirical Datasets

In comparisons of DEC and DEC* on empirical datasets, likelihood was always better under DEC*. Thus, AIC always selected DEC* as the better model over DEC (as the two models have the same number of parameters). In 10 out of 15 empirical datasets, AIC selected DEC* over DEC+J (Supplementary Table 1). DEC+J has an extra parameter relative to DEC*, so if likelihoods were equal between DEC+J and DEC*, DEC* would be preferred by AIC. However, in all but one of the cases where AIC preferred DEC* over DEC+J, the likelihood was itself better with DEC* (which is possible, as the two models are not nested). The exception was *Psychotria*, where AIC gives DEC* 50.2% of the model weight despite slightly higher likelihood for DEC+J (model weight 20.4%).

Unlike the simulated data, for over half the empirical datasets the extinction rate inferred by DEC was substantially higher than zero, ranging from 16% to 546% of the estimated value of the dispersal rate. For DEC*, the extinction rates were even higher relative to dispersal: only for one empirical dataset was the extinction rate indistinguishable from zero, for the rest the extinction rate was between 3.2 and 1389-fold higher than dispersal rate (median 104-fold higher). In some cases, the estimated extinction rate was at the maximum allowed by the program; modifying it to increase the bound by tenfold improved the likelihood by a median of 0.061 log likelihood units and increased the extinction estimate up to the new maximum in most cases. The small magnitude of improvement, with the large magnitude of change in the estimate, suggests that the likelihood surface is very flat but that the unconstrained maximum likelihood estimate would be even higher. More simply put, for these datasets, the best estimate of extinction is extremely high, which would mean that after a species expands its range it nearly instantly contracts it (into either the new region or back to the old region). In only three of nine of these datasets was DEC+J chosen over DEC*, despite the apparent evidence for a jump-like dispersal model.

### Model Adequacy

In addition to model choice, a key question to examine with new models is model adequacy: how well does the model fit overall? Even the best-fitting model may not do a good job predicting the data, which would point to the need for new models to better match reality. This has been increasingly emphasized in phylogenetics (Goldman 1993; Bollback 2002; O’Meara 2012; Beaulieu et al. 2013b; Pennell et al. 2015). To see if the DEC* model adequately describes the data, we counted the number of occupied areas estimated for each node and compared this between DEC and DEC* for each empirical dataset. We work under the assumption that the present should look like the past: a clade of island endemics is more likely to have been island endemics for much of their history, rather than being composed of very widespread species that only at the present suddenly became endemic to single islands. Of course, there are processes that could make the present not resemble the past (i.e., a sudden change in climate causing suitable habitat to be divided into isolated patches), but this assumption should hold in most groups. For all but two empirical datasets, the DEC* model was the more adequate model, with estimated range sizes at ancestral nodes more closely matching the estimated the mean range sizes observed at the tips of the phylogeny (Fig. 3). Inference under DEC usually yields ancestral distributions that are very widespread, which is not the case under DEC* (Fig. 3 and Fig. 4).

**Figure 3.**
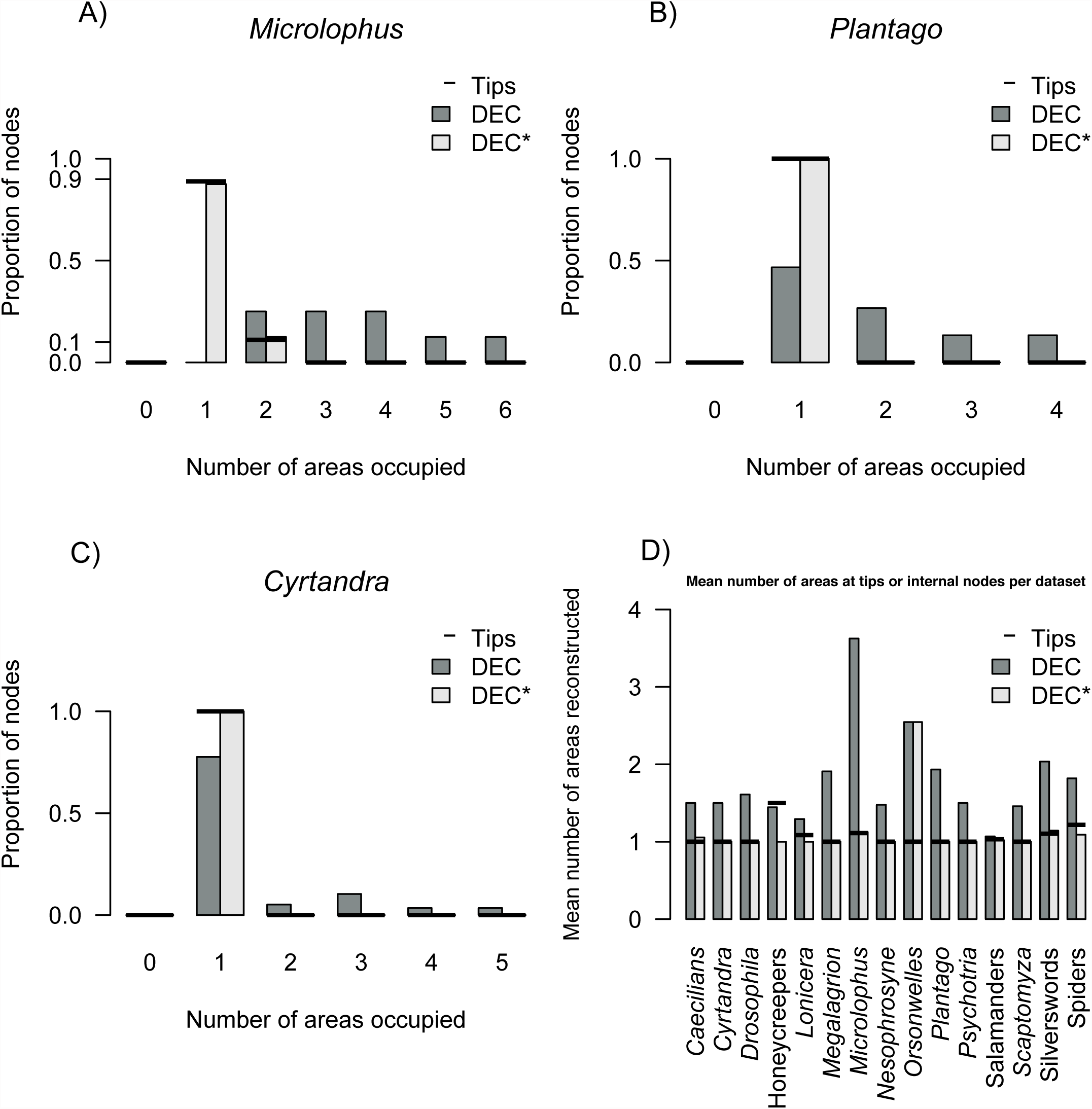
Model adequacy plots based on the number of areas occupied at nodes for the empirical datasets of the Galapagan *Microlophus* (A), Hawaiian *Plantago* (B), Pacific *Cyrtandra* (C) reconstructed with DEC or DEC* versus the tips which represent the current number of areas for each group. In each empirical case (A, B, C), DEC* was able to ancestrally infer the same number of areas as the tips in the current range. DEC inferred ranges that were more widespread. The last plot depicts the mean number of current areas at tips versus the mean number of areas estimated at nodes through DEC or DEC* for each group (D). In each case except for two, DEC* was able to estimated the ancestors to occupy the same number of areas as the tips.

**Figure 4.**
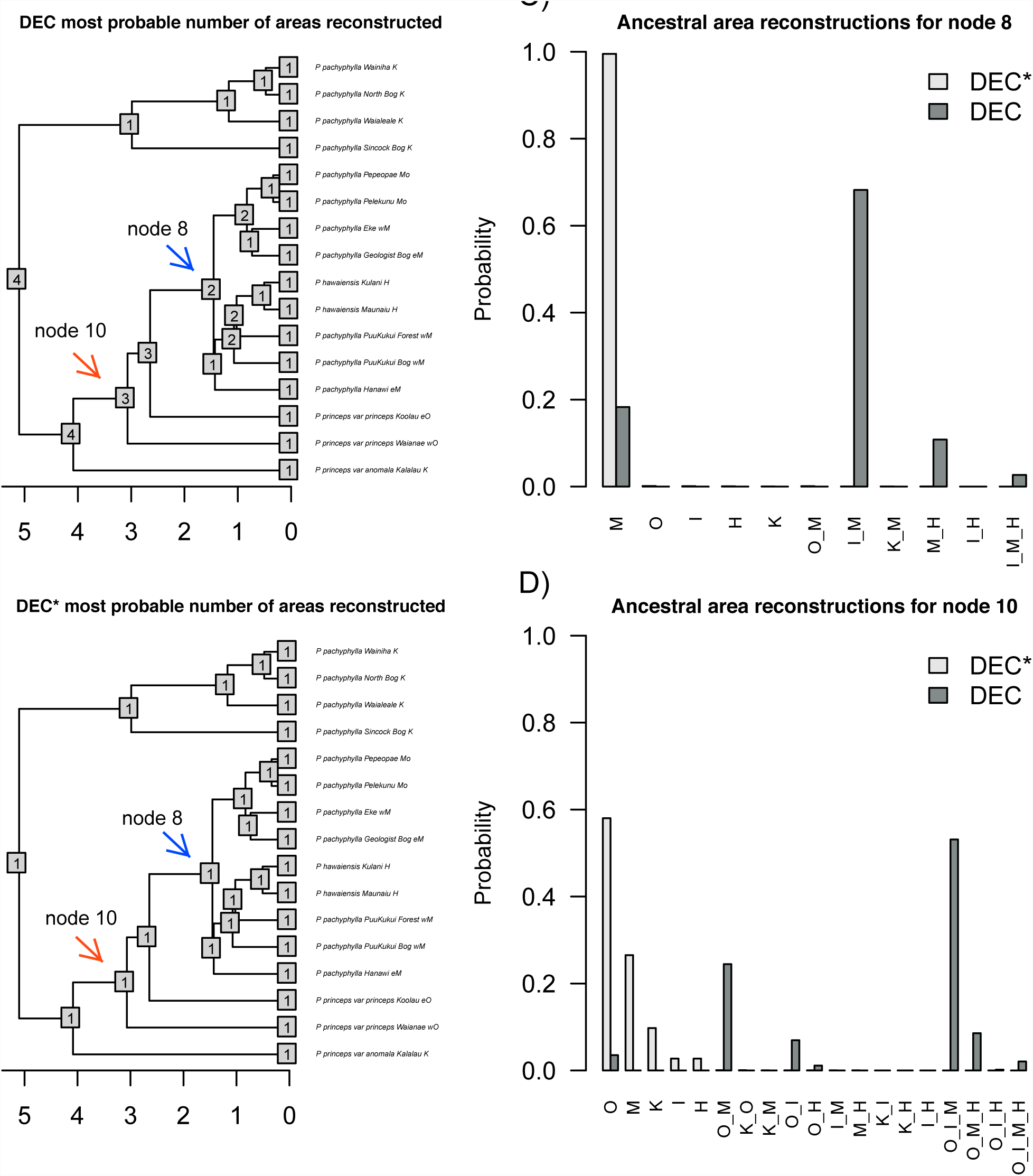
Most probable number of biogeographic areas estimated in the *Plantago* clade under the DEC (A) versus the DEC* model (B). The number of areas at tips is also shown. The estimated ancestral range probabilities under DEC versus DEC* for node 8 (C) and node 10 (D) are shown. Ancestral state estimates under the DEC model are more widespread than under DEC*, therefore providing a different biogeographic history (C and D). Nodes closer to the root provide more variance in the probable ranges estimated (D).

## Discussion

Given the results, we argue that DEC* should be considered for use in biogeographic models. Testing models should be an intrinsic part of the research process, so most users should try DEC, DEC*, DEC+J, and future models, but if one were limited to just one model, in most cases DEC* would be preferred, based on the empirical results presented here. DEC* does a more adequate job at estimating ancestral ranges than does the canonical DEC model. However, while the median extinction and dispersal parameters were better estimated under DEC* than with DEC, the RMSE of the estimates on a linear scale was better under DEC, and DEC* often returns very high estimates for extinction rate. For estimating ancestral areas, DEC* is probably the better model, but we urge extreme caution when treating its rate parameters as parameters of interest rather than nuisance parameters. Of course, ancestral states are known to be difficult to estimate well (Cunningham 1999; Oakley and Cunningham 2000), so biologists should expect a great deal of uncertainty with estimates. We also note that the estimates of uncertainty in this model are always underestimates, due to other uncertainty (topology, branch lengths, states) that is typically not accounted for.

Another caveat to the use of DEC* is its treatment of the phylogeny: it assumes range evolves on a tree but that biogeography does not directly influence speciation or extinction. Speciation often seems influenced by geographic context (Mayr 1963), such as through the divergence of two isolated populations. While DEC, DEC*, and DEC+J allow subdivision of ranges at speciation events, these models do not, for example, fit a higher speciation rate to species with larger ranges. There are models that jointly fit the diversification process and process of biogeographic evolution, such as GeoSSE (Goldberg et al. 2011) and ClaSSE (Goldberg and Igić 2012). However, even though these models are more realistic, they can require rather large data sets (Davis et al. 2013) and are feasible only for very few areas. The empirical datasets used in biogeographic models are typically small in comparison; the ones used in this study had an average of 75 taxa and 5.53 areas from all datasets, and a range of 4 to 10 areas and 9 to 469 taxa. Recent work (Matzke 2014b) showed that DEC vs. DEC+J model choice appears robust to some commonly-postulated SSE processes (speciation and extinction depending on range size), and that ancestral range estimation is reasonably accurate if model choice is performed and if the dispersal rate is low, suggesting that for datasets that limit the power of SSE models, the DEC* model can still be used, with caution.

It is important to emphasize that the DEC* model is still relatively simple. Though some complexity can be incorporated with different dispersal rates between areas or at different time points, all species are treated as having the same rates of dispersal and extinction at a given time. We know, however, that species in a clade may vary in traits affecting successful dispersal (ability to inbreed, resting stages, wind versus animal dispersal, tolerance of saltwater, and so forth) or extinction (body size, trophic level, thermal tolerance, and so forth) and this variation is not yet incorporated in any of these models. There are additional sources of heterogeneity that also may result in misleading results if not incorporated.

The very high extinction estimates by DEC*, especially in empirical datasets, was unexpected. One partial explanation may come from the empirical distribution of range sizes in the empirical studies presented here, where the vast majority of species were found in just one area. Under DEC* (or DEC) the only way for a species to change its area is through expansion to a new area (i.e., dispersal), followed by other events. Given that all the studies have species in different areas (there is little point to inferring biogeographic history for a clade that only ever occupies one area), there must be a nonzero dispersal rate. Lineages can reduce range size in two ways: at cladogenesis, or through range contraction along branches. However, when *e* is high with respect to *d* and the speciation rate, lineages will spend almost no time in widespread ranges. Therefore, widespread ranges are essentially never available at cladogenesis, and all speciation will be sympatric. Moreover, on terminal branches, which represent over half the branches on the tree, any necessary dispersals cannot be “undone” by contraction at cladogenesis, and so the only way to have some dispersals along terminal branches, but have observed species in just one area each, is to have a substantial extinction rate. DEC* with high *e* is a model with all range-change effectively occurring in anagenetic “jumps” along the branches: any expansion is followed almost immediately by a contraction. The fact that DEC* often outperforms DEC+J probably indicates that in these cases, the probability of sister taxa living in different areas correlates better with the branch length between the taxa than with the number of speciation events recorded in the observed tree. We expect a DEC*+J model may incorporate the best of both of these models, and that adding “*” to other models may also be beneficial.

The major use of parametric models in historical biogeography is for ancestral state estimation. For model adequacy, we expect that the present tends to resemble the past, so a model where past distributions are similar to present ones is probably a better fit to the data. For empirical datasets used in biogeography, tip taxa are most often in one area. But, DEC often estimates ancestors as being in many areas, because the DEC model allows a transition into the null range, and since null ranges are not observed, the inference is pushed towards a low extinction rate. In contrast, the DEC* model returns estimates at internal nodes that usually resemble the number of areas present in the tips (see Fig. 4). DEC* may return nearly equally likely single areas rather than a more confident estimation of the ancestral state being a union of areas. In many cases, especially given observed species that occupy individually few areas, this uncertainty about which single area an ancestor occupied represents reality. However, even though uncertainty exists in ancestral range estimates, statistical model choice is a fruitful way to assess models against the data.

## Acknowledgements

We thank the HOFF Lab Group, James Fordyce, Daniel Simberloff, and Sally Horn for helpful discussions. Support for KAM has been provided by the National Institutes of Health Program for Excellence and Equity in Research Grant [R25 5R25GM086761-06] at the University of Tennessee and from the University of Tennessee, Knoxville Ecology and Evolutionary Biology Department Summer Funding and Chancellor’s Funds. JMB and NJM were by the National Institute for Mathematical and Biological Synthesis, an Institute sponsored by the National Science Foundation, the U.S. Department of Homeland Security, and the U.S. Department of Agriculture through NSF Award #EF-0832858, with additional support from The University of Tennessee, Knoxville.

